# Rationalizing PROTAC-mediated ternary complex formation using Rosetta

**DOI:** 10.1101/2020.05.27.119347

**Authors:** Nan Bai, Palani Kirubakaran, John Karanicolas

## Abstract

PROTACs are molecules that combine a target-binding warhead with an E3 ligase-recruiting moiety; by drawing the target protein into a ternary complex with the E3 ligase, PROTACs induce target protein degradation. While PROTACs hold exciting potential as chemical probes and as therapeutic agents, development of a PROTAC typically requires synthesis of numerous analogs to thoroughly explore variations on the chemical linker; without extensive trial and error, it is unclear how to link the two protein-recruiting moieties to promote formation of a productive ternary complex. Here, we describe a structure-based computational method for evaluating suitability of a given linker for ternary complex formation. Our method uses Rosetta to dock the protein components, then builds the PROTAC from its component fragments into each binding mode; complete models of the ternary complex are then refined. We apply this approach to retrospectively evaluate multiple PROTACs from the literature, spanning diverse target proteins. We find that modeling ternary complex formation is sufficient to explain both activity and selectivity reported for these PROTACs, implying that other cellular factors are not key determinants of activity in these cases. We further find that interpreting PROTAC activity is best approached using an ensemble of structures of the ternary complex rather than a single static conformation, and that members of a structurally-conserved protein family can be recruited by the same PROTAC through vastly different binding modes. To encourage adoption of these methods and promote further analyses, we disseminate both the computational methods and the models of ternary complexes.

**Significance Statement:** Recent years have brought a flood of interest in developing compounds that selectively degrade protein targets in cells, as exemplified by PROTACs. Fully realizing the promise of PROTACs to transform chemical biology by delivering degraders of diverse and undruggable protein targets has been impeded, however, by the fact that designing a suitable chemical linker between the functional moieties requires extensive trial and error. Here, we describe a structure-based computational method to predict PROTAC activity. We envision that this approach will allow design and optimization of PROTACs for efficient target degradation, selection of E3 ligases best suited for pairing with a given target protein, and understanding the basis by which PROTACs can exhibit different target selectivity than their component warheads.

## Introduction

PROteolysis TArgeting Chimaeras (PROTACs) are heterobifunctional small molecules containing two functional moieties: a targeting ligand (warhead) for the protein of interest, and a ligand that recruits an E3 ubiquitin ligase. If these two groups are connected via a carefully-chosen chemical linker, the resulting compound can induce formation of a ternary complex that includes the target protein, the PROTAC, and the E3 ligase. The resulting proximity between target protein and E3 ligase can then lead to ubiquitination of the target protein, and ultimately its degradation [1].

In essence, PROTACs represent a chemical biology approach for extending the natural activity of RING-type E3 ligases. This most-populous subclass of E3 ligases act as adaptors, by simultaneously binding to their target protein and a ubiquitin-charged E2 ligase: this enables the E2 ligase to transfer ubiquitin directly to the substrate protein [2]. The E2 ligase appears not to encode its own substrate selectivity, but rather relies completely on the E3 ligase to recruit appropriate substrate proteins; it has long been known that this makes it susceptible to hijacking by viral proteins that redirect E2 ligases for degrading host proteins involved in defense against pathogens [3]. Small molecules can also redirect E3 ligase activity in this manner: the most notable of these is thalidomide, which (inadvertently and tragically) binds to the E3 ligase cereblon (CRBN) and redirects it to degrade certain key neosubstrates [4-7].

The perspective that PROTACs act by “retrofitting” an E3 ligase with new substrate selectivity makes immediately clear the many strengths of this approach relative to classical competitive inhibitors. Principal among these advantages are the fact that PROTACs need not engage the target protein’s active site (thus expanding the realm of the druggable proteome) [8], and the fact that PROTACs can exhibit catalytic turnover (thus allowing degradation of the target protein at a substoichiometric ratio of PROTAC) [9].

Over the past decade, PROTACs have been developed to address an ever-expanding collection of target proteins. Owing to the availability of well-validated chemical matter to repurpose as PROTAC warheads, the majority of early reports focused on degraders of hormone receptors [10-13], BET-family proteins [14-16] and kinases [17-33]. More recently, the applicability of PROTACs has been broadened to include multi-functional proteins (e.g., TRIM24 [34], SMARCA2 [35] tau [36,37]), and HaloTag7-fused proteins for chemical genetics [38]. In parallel, the space of E3 ligases used has also expanded: whereas early studies focused exclusively on CRBN and Von Hippel-Lindau protein (VHL), more recent work has used E3 ligases MDM2 [39], IAP [40], RNF4 [41], βTRCP [42], parkin [42], and DCAF16 [43]. Collectively, there appear to be an apparently endless set of intriguing biological targets and a large set of E3 ligases that might be used to selectively degrade them.

A key challenge in this rapidly-growing field, however, remains unresolved: how to design effective PROTACs in a rational manner. A important early paper in this field advocated for rapidly varying both the target-engaging warhead and the E3-recruiting moiety in search of a useful starting point [17], and this strategy is still used today. This approach can only hope to capture only a fraction of the candidate PROTACs one might envision for degrading a given target, however, because the linker used to join these groups also plays a critical role in efficacy. Since the earliest days of PROTAC design, the dominant strategy has simply been to synthesize and test many tens – or hundreds – of variants in which the length and chemical composition of the linker are systematically varied [44,45]. Indeed, given sufficient perseverance with respect to conjugation patterns and linker length/composition, even target/ligase pairings that initially seemed unsuitable have been optimized into potent and selective degraders [46].

Here, we sought to develop a structure-based modeling approach for predicting the efficacy of a given PROTAC; if successful, we envisioned that this approach could be used for optimization of activity without needing to explicitly synthesize and test numerous analogs. Our study seeks to do so by modeling the PROTAC’s ability to induce formation of the target protein/PROTAC/E3 ligase ternary complex. It has been shown that a PROTAC’s ability to bind the target protein is not the primary determinant of whether efficient degradation occurs; rather, it has been suggested that formation of the ternary complex is limiting (i.e., the ability of the PROTAC to simultaneously bind both the target protein and the E3 ligase) [20,22]. It has further been shown that PROTAC selectivity does not necessarily mimic that of its target-binding warhead, but rather depends sensitively on the linker [20,23,24,33]: accordingly, we therefore additionally explore whether the same structure-based modeling can be used to rationalize how and why PROTAC selectivity differs from that of the warhead.

To encourage adoption of these computational methods and to facilitate additional analyses of the results presented below, both the computational methods and the models of the ternary complexes described herein are disseminated with this work (see *Methods*).

## Computational Approach

A PROTAC can be naturally decomposed into its three component parts: the warhead that binds the target protein, the E3 ligase ligand, and the linker. PROTACs are most often built upon well-characterized warheads, and thus the structure of the warhead in complex with its target protein is typically known *a priori*. Crystal structures of many widely-used E3-recruiting ligands have also been solved in complex with their cognate E3 ligases [47,48]. The interactions of each component within its respective binding site is unlikely to change in the context of the PROTAC ternary complex; from a structural standpoint then, the key challenge entails determining the placement of the two proteins relative to one another.

This challenge can be viewed from two perspectives. From the vantage point of the PROTAC, if the linker conformation is known ahead of time (or is sampled thoroughly enough), the ternary complex can be built by alignment of each protein back onto the moiety responsible for recruiting it [30]. From the vantage point of the protein pair, conversely, the ternary complex can be constructed by building a linker conformation to connect the two protein-recruiting moieties. Both strategies were compared in a recent study [49]; due to sampling considerations, this work (as well as our own early experimentation) found superior performance building linkers into pre-placed protein pairs (i.e., the latter approach).

The observation that many effective PROTACs exhibit cooperativity in assembling the components of the ternary complex [35,50,51] implies the presence of adventitious favorable interactions between the target protein and the E3 ligase in the ternary complex. Our overarching strategy was therefore to use protein-protein docking to build up a collection of candidate binding modes with some degree of complementarity, and then use structural and energetic refinement to filter for those consistent with the specific linker present in the PROTAC. Protein-protein docking has previously been used in a similar manner to compile likely poses for PROTAC ternary complexes [52], albeit without explicitly building the bridging linker into these models.

Below we summarize our computational pipeline for building models of PROTAC ternary complexes. All code is implemented in the Rosetta software suite [53]; complete details, including command lines for running these calculations, are included in the *Methods* section.

### Step 1: Building diverse binding modes

Initial candidate binding modes are generated via protein-protein docking in Rosetta, using as a starting point the structures of each protein (the protein targeted for degradation and the E3 ligase) bound to their cognate ligands (the warhead and the E3-recruiting moiety, respectively). Because formation of the ternary complex requires an orientation in which the two proteins position their respective ligands close to one another, this in turn restricts the space of suitable docked models. Accordingly, we elected to re-purpose a docking protocol originally developed for antibody-antigen modeling, that was built for scenarios in which the approximate surface patches involved in binding are known (i.e., an interaction between a known epitope on the antigen and the antibody’s CDR loops) [54].

Drawing from standard practice for this docking pipeline, we begin by manually placing the two proteins near one another, with the ligands facing each other. Docking is carried out by moving one protein with respect to the other, while keeping fixed all internal degrees of freedom for both proteins. Each protein’s bound ligand is included in the simulation and moves with the protein, such that the protein-ligand interactions are not disturbed. A total of 50,000 bound poses are generated from independent docking trajectories: based on the move set used in docking (see *Methods*), the resulting collection of models includes very diverse binding modes that thoroughly capture potential arrangements of the two proteins (**Figure 1a**). The resulting models are then ranked on the basis of the protein-protein interaction energy (including contributions from the bound ligands); the top 10% of these are advanced for further refinement.

**Figure 1:**
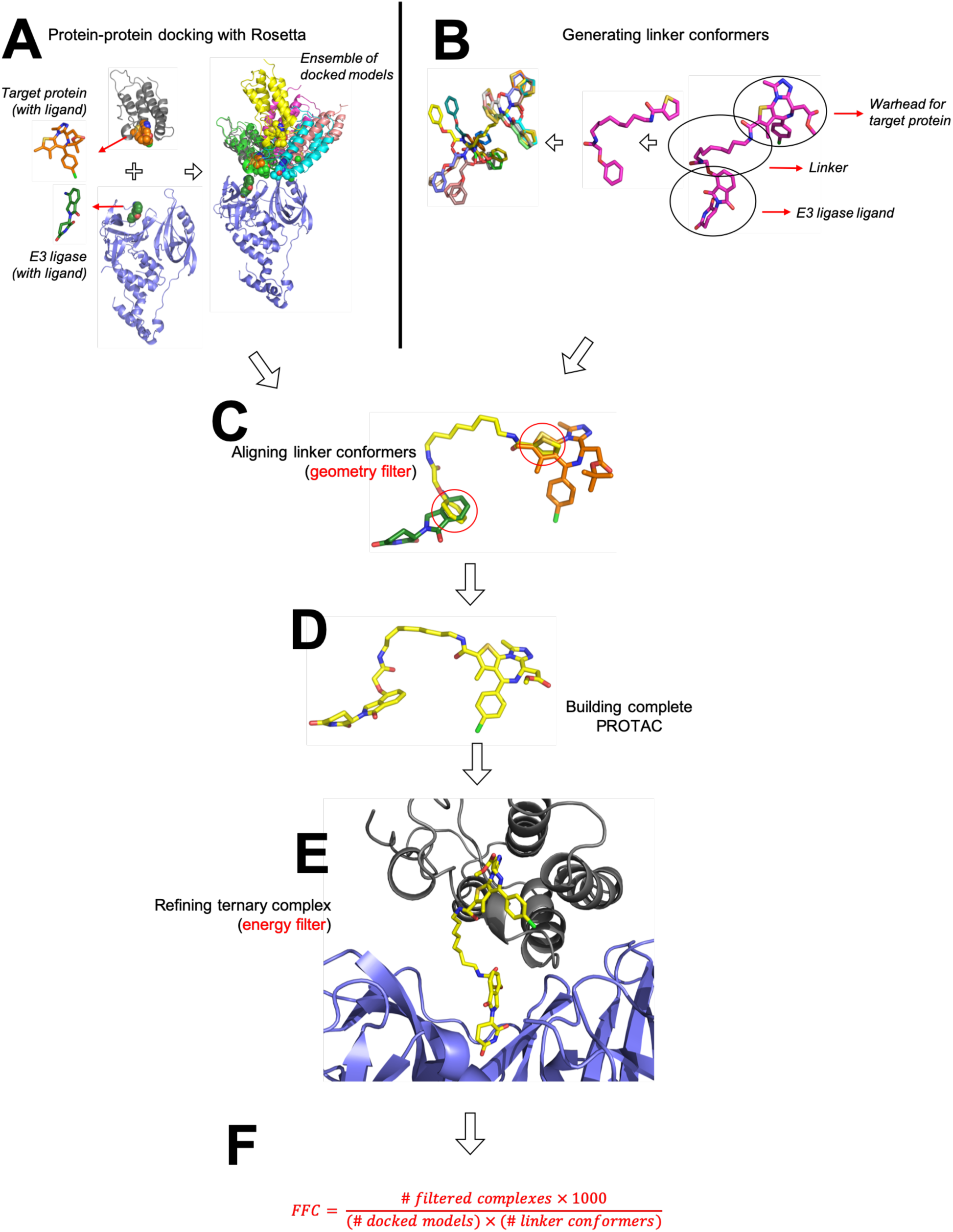
Overview of the computational approach. **(A)** The target protein and the E3 ligase are first docked using Rosetta. Both the target protein and the E3 ligase harbor a bound ligand at this stage (the PROTAC’s warhead and its E3-recruiting group, respectively); this is used to restrict docking to binding modes in which the ligands are close to one another. The resulting collection of docked models does not include any information about any particular linker, and can therefore be re-used to explore and compare multiple linkers. **(B)** Candidate linker conformations are generated using OMEGA. Conformers include “stubs” either end that correspond to the attachment points on the two protein-recruiting moieties. **(C)** For each docked model, the set of linker conformations is screened to find those that place the “stubs” back onto their corresponding atoms in the docked model; this allows rapid identification of low-energy linker geometries compatible with the docked binding mode. **(D)** When a match is found, the complete PROTAC molecule can be trivially re-built from the component parts. **(E)** Because the protein-recruiting moieties of the PROTAC have not moved in building the complete PROTAC, the PROTAC’s position relative to the protein components is also known: this allows for trivial assembly of a complete ternary complex model, which is then subjected to energy minimization in Rosetta. **(F)** After filtering, the *<u>F</u>*raction of *<u>F</u>*ully-compatible *<u>C</u>*omplexes (*FFC*) is calculated. This measures how well a given PROTAC linker is geometrically and energetically compatible with the preferred interaction modes of the protein pair.

### Step 2: Generating candidate linker conformations

Independently from this docking step, we assemble a collection of low-energy linker geometries. Capturing the breadth of viable linker conformations is much akin to modeling the flexibility of drug-like molecules when docking them to proteins. Accordingly, we use the OMEGA software [55,56] to compile a set of “conformers” (conformational isomers) corresponding to the PROTAC linker. The modeled linker also contains on either end a chemical “stub” corresponding to several atoms of the functional group to which it is attached in the PROTAC (i.e., part of the warhead or the E3-recruiting moiety) (**Figure 1b**).

OMEGA uses a rule-based search strategy to build starting structures, coupled with molecular mechanics minimization; a collection of conformers is built up by requiring that each new conformer is not the same as one that has previously been generated. Our goal at this stage is to thoroughly sample the linker region’s internal degrees of freedom. We therefore continue generating new OMEGA conformers either until we accumulate 1000 non-redundant structures, or until sampling converges with fewer than 1000 structures.

### Step 3: Building initial models of the ternary complex

Having compiled a collection of candidate binding modes in Step 1, we next filter these on the basis of whether their bound ligands (the warhead and the E3-recruiting moiety) can be bridged by one of the low-energy linker conformations from Step 2. In essence, this requirement captures the crux of designing a PROTAC: selecting a linker compatible with a reasonable arrangement of the two proteins, in order to induce formation of a ternary complex. The collection of docked structures from Step 1 was built without any specific PROTAC in mind; it is therefore only now at this filtering step that the space of allowed binding modes is restricted based on the linker present in a particular PROTAC.

To filter the collection of docked structures, we make use of the chemical “stubs” present in the conformers built in Step 2. These particular atoms are present both in the docked protein structures, and also in the linker conformers. To gauge whether a specific linker conformation would suitably span the ligands in a given binding mode, we structurally align these shared atoms and evaluate the RMSD (root mean square deviation) between their positions (**Figure 1c**). Through preliminary exploration we found that RMSD values less than 0.4 Å yielded binding modes suitable for further refinement, and thus we adopted this as a cutoff to determine which models to pass along for further refinement. As will be demonstrated below in the *Results* section, the requirement that the component parts of the PROTAC can be connected by the linker represents a very stringent filter that eliminates the vast majority of docked models.

For each combination of docked binding mode / linker conformer that yields low-RMSD overlap of the shared stub atoms, we proceed to build a complete model of the ternary complex. To do so, we simply merge the aligned linker conformer with the coordinates of the bound ligands: this step provides coordinates for the complete PROTAC molecule (**Figure 1d**). The resulting model of the ternary complex is then subjected to energy minimization using the Rosetta energy function [57], which eliminates slight distortions in bond geometries as well as potential steric clashes between the linker and the protein components (**Figure 1e**).

### Step 4: Analyzing ternary complex models

For each of the refined models of the ternary complex from Step 3, we then evaluate the interaction energy amongst all components of the system: this entails summing the interaction energies of the PROTAC with each of the two proteins, and the interaction energy of the proteins with each other. To normalize these energies in a meaningful way, we first determine the median interaction energies for the original complete set of original docked models for this protein pair (i.e., in the absence of a PROTAC linker), and set this as a cutoff value. We then filter the fully-refined models of the ternary complex, keeping only with an interaction energy better than this cutoff value.

In essence, passing this filter indicates either that a given model of the ternary complex utilizes a better-than-average binding mode available to this pair of proteins, or else that the linker itself contributes favorably to the interaction. Conversely, this filtering step is designed to remove any models in which the protein-protein interaction is suboptimal, the PROTAC linker clashes with one of the other components in the system, or one of the protein-recruiting moieties of the PROTAC has been disturbed from its initial positioning in the cognate protein’s binding site.

After this final filtering step, we report the *<u>F</u>*raction of *<u>F</u>*ully-compatible *<u>C</u>*omplexes (*FFC*): essentially, the fraction of the initial docked models that emerge after this extensive geometric and energetic filtering, relative to the original (linker-independent) set of docked models (**Figure 1f**). Ultimately, *FFC* is designed to provide a measure of how well the restrictions from the PROTAC align with the inherent preferences found in the original ensemble of docked models.

## Results

While there have been several published crystal structures of ternary complexes [35,50,52], there are not yet enough to allow for a thorough and rigorous evaluation of our method. Further, it remains unclear whether a single static snapshot is sufficient to understand the structural basis for PROTAC activity. For these reasons, we instead sought to evaluate the utility of our computational approach by comparing to the extent of target protein degradation induced in cells treated with the PROTAC. This is by far the most common form of data available in literature describing PROTAC development, allowing for multiple interesting points of comparison.

That said, our computational approach models only the ternary complex, and neglects other factors that may influence observed cellular activity; these factors are numerous, and surely include the PROTAC’s cell penetrance, efflux, kinetics of ternary complex formation, ubiquitination efficiency once the ternary complex is formed, and rate of proteasomal degradation of the ubiquitinated target protein [58]. While techniques have been developed to individually probe each of these steps, as well as methods for probing ternary complex formation in live cells [59], results have not been reported in a sufficient number of cases to allow careful benchmarking. Instead, here we seek to test the hypothesis that ternary complex formation alone – which we explicitly model – can serve as a suitable proxy for predicting cellular activity (degradation of the target protein).

To validate this approach, we compiled data from literature describing optimization and characterization of PROTACs addressing diverse target proteins using multiple E3 ligases. Due to the availability of complete data across chemical series, we elected to focus on CRBN-recruiting degraders of Brd4^BD1^ [52,60] and CRBN/VHL-recruiting degraders of c-Met/EphA2/STK10/p38α/p38δ [20]. As described in detail below, the first of these case studies demonstrates that our computational method can rationalize the activity of related PROTACs in response to different linker lengths; the second case study shows that these models not only recapitulate the selectivity of a given PROTAC with different (kinase) targets, but that they also recapitulate efficacy of PROTACs built to recruit different E3 ligases.

### Dependence of PROTAC activity on linker length

PROTAC design to date has inevitably required synthesis and testing of numerous analogs with slightly-varied linkers, to optimize degradation of the target protein; as a starting point for evaluating our method, therefore we asked whether computational design could have facilitated this laborious process. In other words, given a set of linkers, we sought to determine whether our computational approach would correctly prioritize which of these yields the best activity in the context of a particular warhead and E3-recruiting moiety.

As a starting point, we examined two independent series of PROTACs that target the BET-family member Brd4^BD1^ for CRBN-driven degradation [52,60]. Both series use derivatives of the Brd4 inhibitor JQ1 [61] as a warhead and both studies report cellular characterization of a series of analogs in which the linker length is systematically lengthened. Despite their shared warhead and E3-recruiting moiety, however, the two groups attach the linker to the warhead in a different way.

To study these systems, we began by generating a single ensemble of 5000 docked models starting from the JQ1-bound crystal structure of Brd4 and the lenalidomide-bound crystal structure of CRBN (see *Methods*). This ensemble corresponds to the lowest-energy 10% of the raw docking output, and is intended to capture the diversity of potential ways in which these two proteins might form adventitious interactions with one another; accordingly, there is no information on the linker length or attachment point of any specific PROTAC at this stage.

From here, we then built conformers that captured the linkers and attachment points for each of the five PROTACs comprising the “ZXH” series [52] (**Table 1**). As noted earlier, this process entails filtering the set of 5000 docked models on basis of their geometric and energetic compatibility with a given linker. We sought to use the number of models that emerge from this extensive filtering as a measure of how efficiently the PROTAC induces ternary complex formation, ultimately with the goal of defining how well this translates to cellular degradation of the target protein.

**Table 1:** Evaluation of the “ZXH” PROTAC series. This series uses JQ1 and pomalidomide to recruit Brd4BD1 for degradation by CRBN. Cellular degradation data are from Nowak et al, 2018.

| PROTAC | Linker | # conformers generated | # geometrically compatible binding modes | FFC | Cellular degradation of Brd4<sup>BD1</sup> (EC<sub>50</sub> in nM) |
| --- | --- | --- | --- | --- | --- |
| ZXH-3-27 |  | 237 | 0 | 0.00 | > 10000 |
| ZXH-2-147 |  | 606 | 21 | 0.01 | 10 - 100 |
| ZXH-2-184 |  | 1000 | 520 | 0.10 | 10 - 100 |
| ZXH-3-26 |  | 1000 | 1214 | 0.24 | 1 - 10 |
| dBET70 |  | 1000 | 1257 | 0.25 | 1 - 10 |

Remarkably, for the five PROTACs comprising the “ZXH” series, the numbers of poses arising from filtering (which we report as FFC, the *<u>F</u>*raction of *<u>F</u>*ully-compatible *<u>C</u>*omplexes) match very closely the trend in experimentally-observed cellular degradation of the target protein (**Table 1**). We observe that linkers with higher FFC values (i.e., the linker is compatible with more of the conformational ensemble from docking) correspond exactly to those with enhanced cellular degradation of Brd4^BD1^, presumably because these PROTACs more effectively induce ternary complex formation in cells. We note that more conformers were generated for the longer linkers, and yet this would not explain the observed trend because FFC is normalized to the number of conformers included in each calculation.

As noted earlier, a separate set of Brd4-degrading PROTACs have been built by attaching pomalidomide to slightly different JQ1 derivative, and using a different attachment strategy [60]. Unfortunately, these compounds were not tested in the same cell lines as the “ZXH” series, which precludes a direct comparison of activities between the PROTACs from these two different studies. That said, the latter study reports a series of very potent degraders [60], and indeed these yield high FFC values when explored through our computational approach (**Table S1**).

**Table S1:**
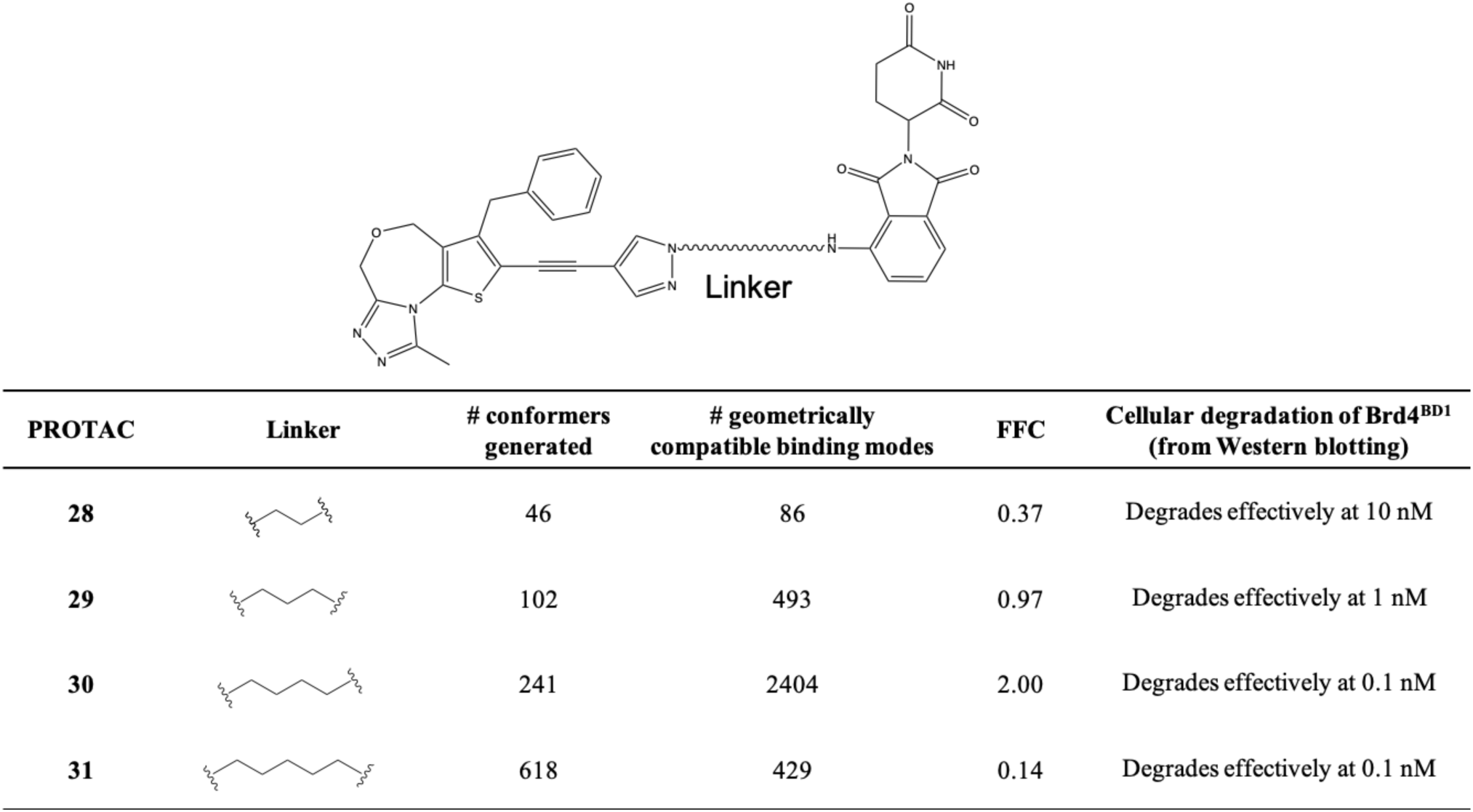
Evaluation of the “28-31” PROTAC series. This series uses JQ1 and pomalidomide to recruit Brd4BD1 for degradation by CRBN. Cellular degradation data are from Qin et al, 2018.

By randomly splitting the original collection of docked models into five subsets, we next evaluated the variance arising from sampling in each of the predicted FFC values for the “ZXH” series of PROTACs: this analysis confirms the statistical significance of the differences between calculated FFC values (see *Methods*) (**Figure 2a**).

**Figure 2:**
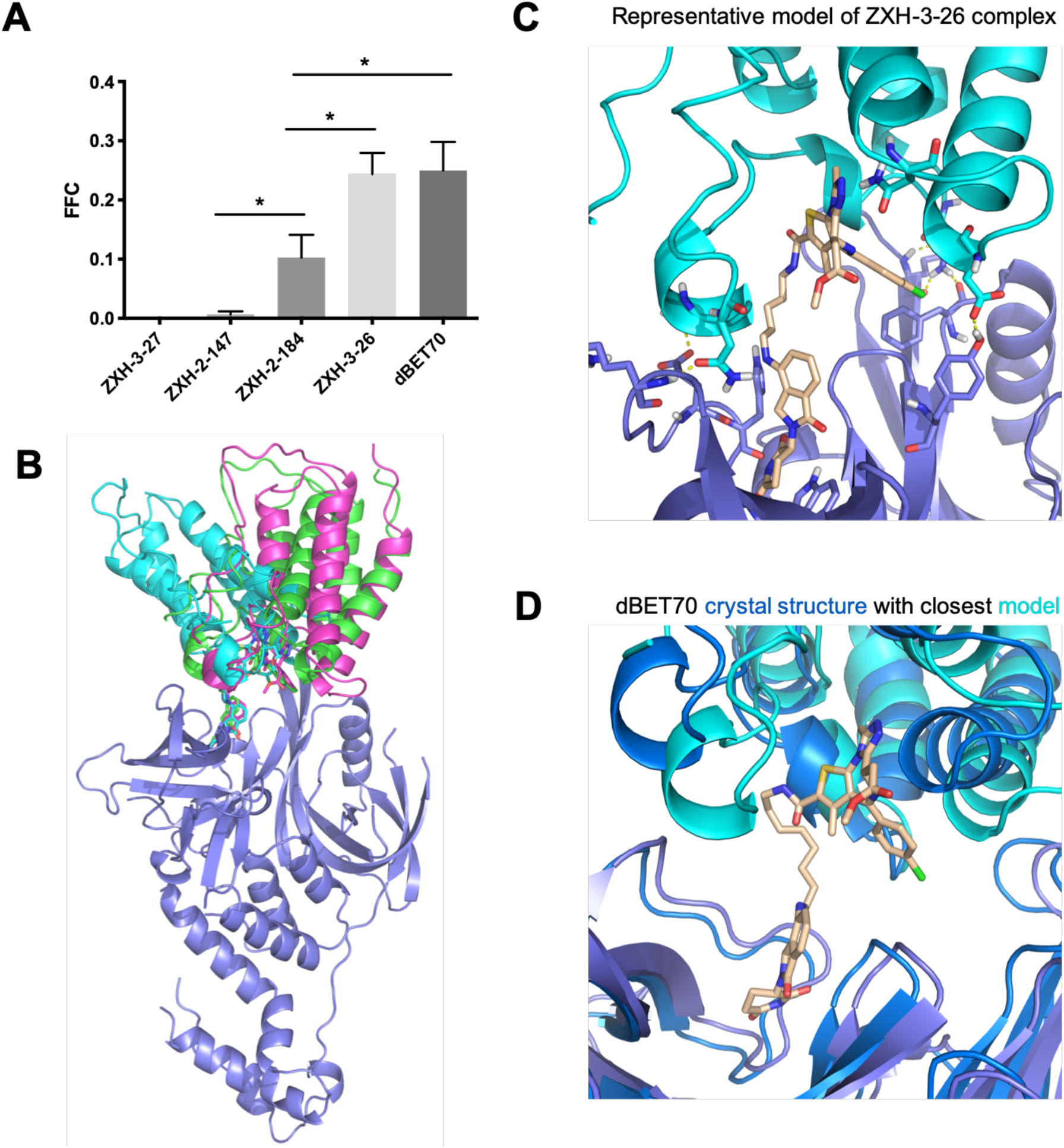
Evaluation of PROTACs that induce CRBN-mediated degradation of Brd4BD1. **(A)** Each PROTAC in this chemical series uses the same warhead and E3-recruiting moiety, allowing a single ensemble of docked models to be used for this collection. Filtering these on geometric and energetic criteria yielded values for FFC (the *<u>F</u>*raction of *<u>F</u>*ully-compatible *<u>C</u>*omplexes) that varied greatly from one linker to another. As shown in **Table 1**, FFC closely matched the trend of relative activity across this chemical series. Error bars correspond to SEM, * indicates p<0.05. **(B)** For each of these PROTACs, the models contributing to FFC do not correspond to a single binding mode. Rather, each PROTAC allows for multiple diverse binding modes that are fully-compatible with the ternary complex. A subset of these models is shown for the PROTAC dBET70. **(C)** A representative ternary complex model from one of the other PROTACs in this series, ZXH-3-26, is shown to illustrate the favorable adventitious interactions found between the component proteins in a representative one of these models. **(D)** The crystal structure of dBET70 was previously solved in complex with BRD4 and CRBN (PDB 6bn9). The collection of ternary complex models generated for dBET70 includes the binding mode observed in this crystal structure. The crystallographic resolution did not allow dBET70 to be unambiguously placed, and therefore the PROTAC is shown only for the modeled structure.

Importantly, we find that the collection of models which pass these stringent filters (i.e., those that contribute to FFC) do *not* represent a single binding mode. Rather, we find that for these PROTACs there are a remarkably diverse set of conformational arrangements allowing for models of the ternary complex that are fully consistent with all geometric and energetic requirements (**Figure 2b**). This observation is consistent with the surprising “plasticity” originally ascribed to the complexes in this series [52]. It is further consistent with the fact that many single-point mutations to either Brd4 or to CRBN (outside of their respective ligand-binding sites) lead to modest changes that enhance or reduce ternary complex formation, without finding mutations that abrogate the interaction completely [52]. A representative model for one such ternary complex model (using the PROTAC ZXH-3-26) shows an example of the adventitious protein-protein interactions observed in these models (**Figure 2c**). Despite the fact that these proteins are presumably not at all evolved to recognize one another, we find that they can exhibit surprising shape complementary along with a number of stabilizing intermolecular hydrogen bonds. The linker itself can also contribute favorably to ternary complex formation in some cases, by making additional interactions with one or both proteins.

The crystal structure for the dBET70 ternary complex has been solved [52], and gratifyingly our collection of models contributing to FFC does include this particular binding mode (**Figure 2d**). Intriguingly, however, the authors surmised even at the time (on the basis of mutational signatures) that this arrangement may not be dominant one in solution, but rather an arrangement that happens to be stabilized by crystal packing. This viewpoint is wholly consistent with our observation that multiple binding modes are likely to contribute to the ternary complexes formed by these PROTACs. Moreover, this example highlights the need to evaluate predicted PROTAC-inducted ternary complex formation against solution (or cellular) observables, rather than use recapitulation of static crystal structures of ternary complexes as a gold standard.

Overall, the results from this exploration of Brd4 degraders lay the groundwork for this computational framework, and demonstrate how modeling of ternary structure complexes could be used to facilitate rational design and optimization of efficacious degraders.

### Dependence of PROTAC activity and selectivity on the choice of E3 ligase

Encouraged by these results, we next applied this computational approach to explore a pair of PROTACs build using the non-selective kinase inhibitor foretinib [62] as a warhead. Using a competitive binding assay, foretinib itself was first shown to inhibit at least a quarter of the human kinome [20]. Addition of a linker with a moiety to recruit either VHL or CRBN reduced foretinib’s inhibition for several of these kinases, but overall both PROTACs retained broad non-selective inhibition. Remarkably, however, cellular treatment revealed that certain kinases were degraded much more efficiently by the VHL-recruiting PROTAC, while others were degraded more efficiently by the CRBN-recruiting PROTAC [20]. Moreover, the extent to which different kinases were degraded was not simply a reflection related to how tightly the PROTAC bound the kinase, since inhibition constants were not correlated with cellular degradation [20]. Given the shared warhead, we hypothesized that these preferences arose due to differences in the ternary complex formation (e.g., in the interactions between the E3 ligase and the kinase); if so, the selectivity of these PROTACs may emerge in our computational modeling approach.

Crystal structures of foretinib in complex with three different kinases have been solved to date (c-Met, EphA2, and STK10) [62,63], and we used all three of these as starting points for our studies of these foretinib-derived PROTACs. Because of interesting differences in PROTAC selectivity between p38α and p38δ, we additionally relied on the close structural conservation amongst inhibitor-bound kinases to build comparative models of foretinib bound to each of these two kinases. Using as a starting point these five foretinib-bound kinase structures (**Figure 3a**), along with the crystal structures of VHL and CRBN bound to their respective recruiting moieties, we then carried out the computational approach described earlier and calculated the Fraction of Fully-compatible Complexes (FFC) for each PROTAC with each kinase (**Table 2**).

**Table 2:**
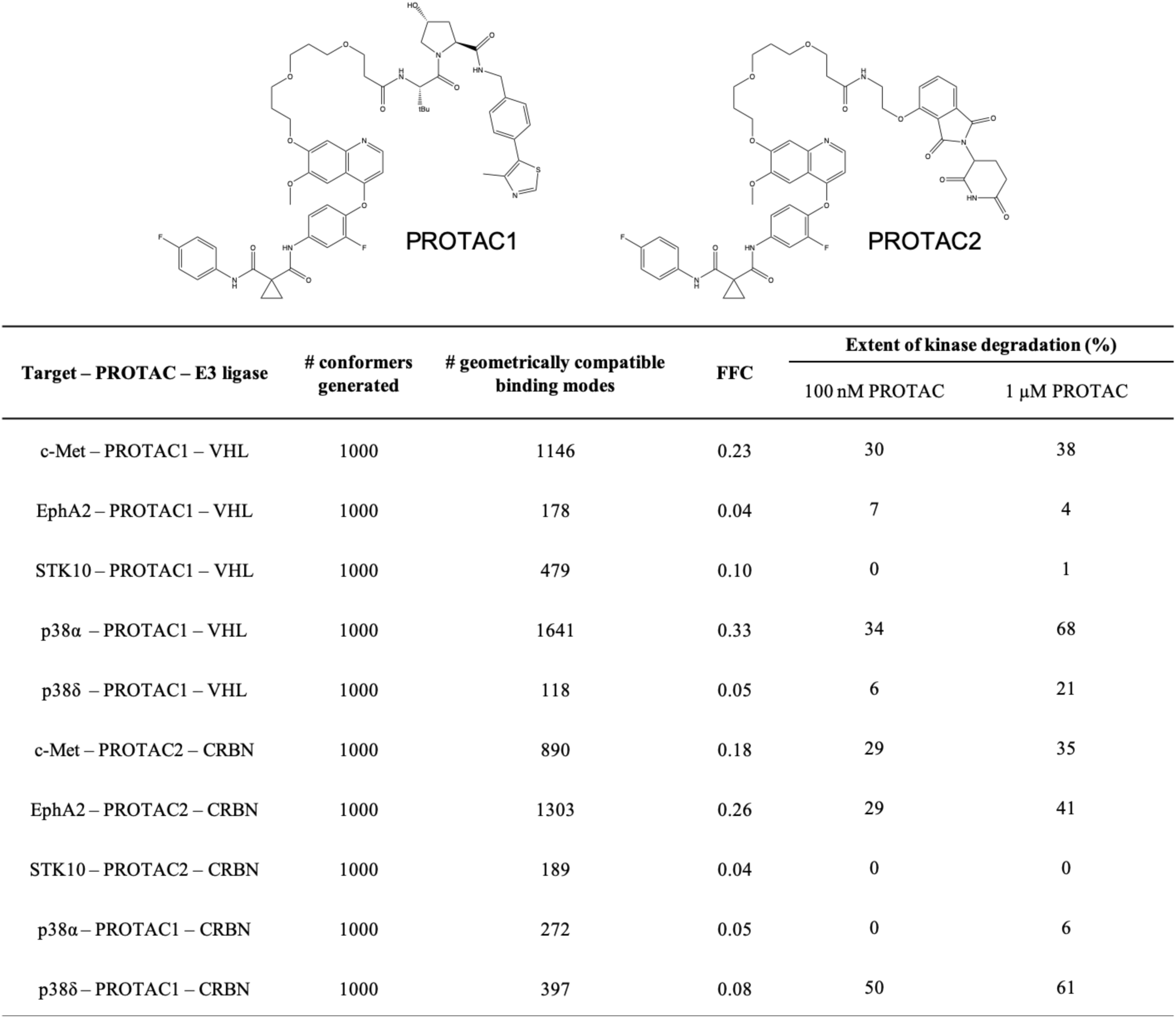
Evaluation of foretinib-derived kinase degraders. These two PROTACs couple non-selective kinase inhibitor foretinib with a moiety to recruit either VHL (PROTAC1) or CRBN (PROTAC2). Cellular degradation data are from Bondeson et al, 2018.

**Figure 3:**
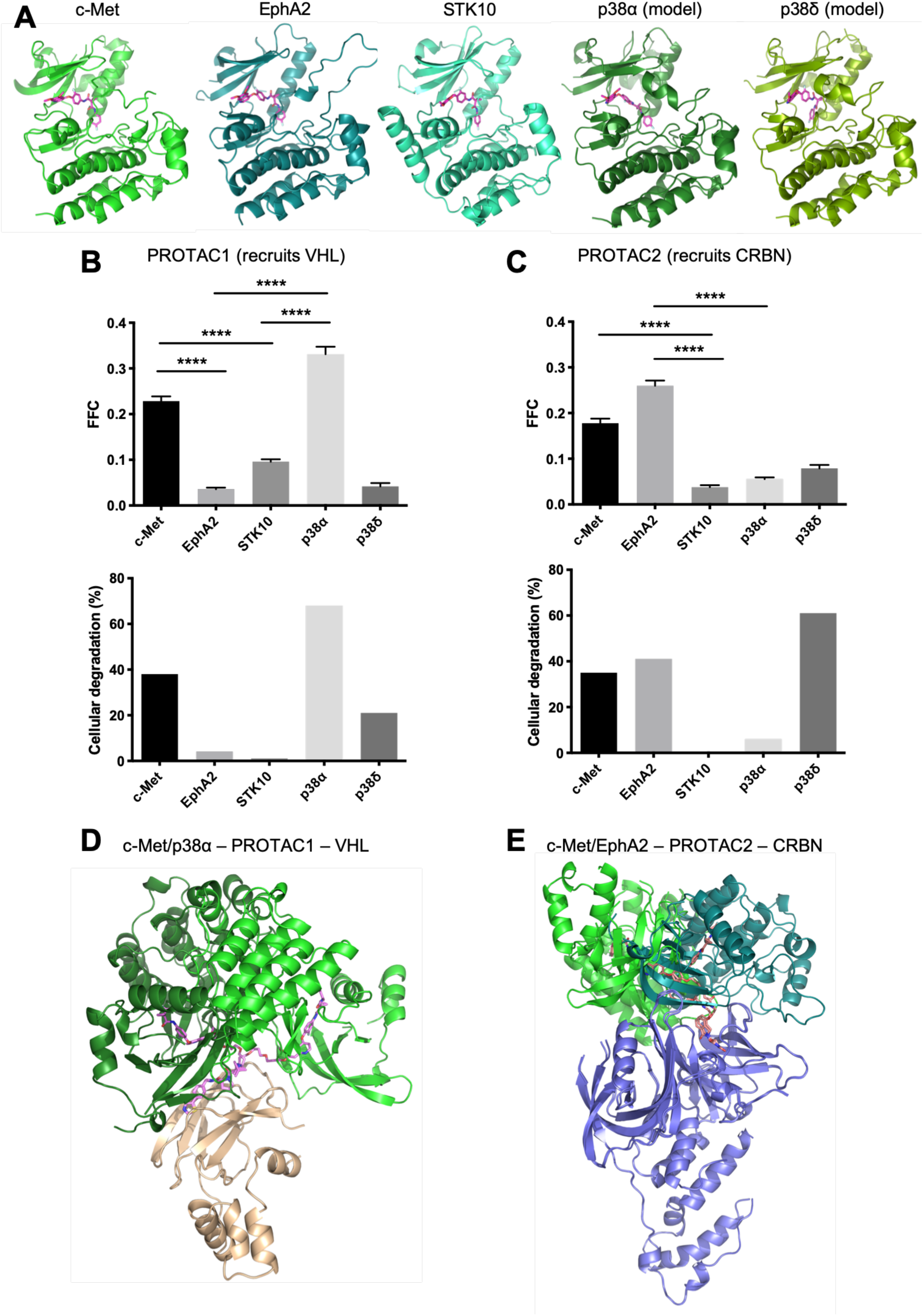
Evaluation of kinase degraders built upon foretinib. **(A)** Calculations were initiated from crystal structures of foretinib in complex with c-Met/EphA2/STK10 or comparative models of foretinib-bound p38α/p38δ. **(B)** For the VHL-recruiting PROTAC1, filtering the resulting models on geometric and energetic criteria yielded values for FFC (the *<u>F</u>*raction of *<u>F</u>*ully-compatible *<u>C</u>*omplexes) that vary amongst these five kinases. The overall selectivity profile is in close agreement with the previously-reported proteomic analysis of kinase cellular degradation induced by this PROTAC. Error bars correspond to SEM, * indicates p<0.05. **(C)** On the basis of FFC, CRBN-recruiting PROTAC2 exhibits a different selectivity profile than PROTAC1. The preferences of PROTAC2 observed in this computational approach also match those observed in the cellular study. **(D)** Upon docking each of these five kinases to VHL (*wheat*), PROTAC1 yields high FFC values for c-Met (*bright green*) and p38α (*dark green*). While the ensembles for both kinases yield multiple fully-consistent binding modes, the preferred binding modes for the two kinases are different from one another. **(E)** Upon docking each of these five kinases to CRBN (*blue*), PROTAC2 yields high FFC values for c-Met (*bright green*) and EphA2 (*turquoise*). Here too, we observe differences in the preferred binding modes for these kinases.

Results from this computational experiment are plotted alongside the corresponding proteomic analysis of cellular degradation reported in the earlier study (**Figure 3bc**). For VHL-recruiting PROTAC1, we observe considerably higher FFC values for c-Met and p38α than for the other three kinases; this is consistent with the extents to which this PROTAC induces their degradation in cells (**Figure 3b**). By contrast, CRBN-recruiting PROTAC2 yields high FFC for c-Met and EphA2: this too is consistent with the cellular effects of treatment with this PROTAC (**Figure 3c**). Collectively then, these results suggest that small differences in the kinase surface alter the extent to which they can be adventitiously recognized by the E3 ligase: by virtue of their different structures, naturally these two E3 ligases exhibit different preferences for the kinases they recognize, which in turn highlights the importance of serendipitously capturing productive protein-protein contacts in PROTAC ternary complexes. As might be expected for such a molecular recognition event, details matter: this is underscored by the fact that strong degradation preferences arise despite the shared global fold of all three kinase domains, as well as the strong similarity between p38α and p38δ.

Given that each of these kinase domains shares a common fold, we anticipated that certain shared features may be present amongst those which were efficiently degraded by a given PROTAC. To our surprise, however, this was emphatically not the case. For PROTAC1, which gave high FFC values for c-Met and p38α, multiple binding modes were observed for both kinases, but the most frequently-observed orientations in the ternary complex were notably kinase-dependent (**Figure 3d**). This observation also held true for PROTAC2, in which both c-Met and EphA2 yielded comparably high FFC values but achieved these values using different binding modes (**Figure 3e**).

In the context of exploring the structural basis for PROTAC1’s preference for p38α over p38δ, a model of the ternary complexes was previously built through a combination of docking and molecular dynamics [20]. This model highlighted the importance of p38α’s Ala40 in the ternary complex, inspiring further experiments that showed mutation of this residue (to arginine) disrupted the ternary complex. Intriguingly, our models also position this residue to make key interactions in the ternary complex, but are otherwise completely different from the previously-proposed models. In particular, the earlier model positions Ala40 near VHL’s Arg69, which is on the remote side of the complex in our models (**Figure S1**). Beyond the fact that we observe ensembles of models that are fully consistent with the geometric and energetic requirements of ternary complex formation (rather than a single static model for each), this observation also underscores the fact that sparse information from mutagenesis experiments are typically insufficient to unambiguously point to a specific binding mode in the ternary complex.

**Figure S1:**
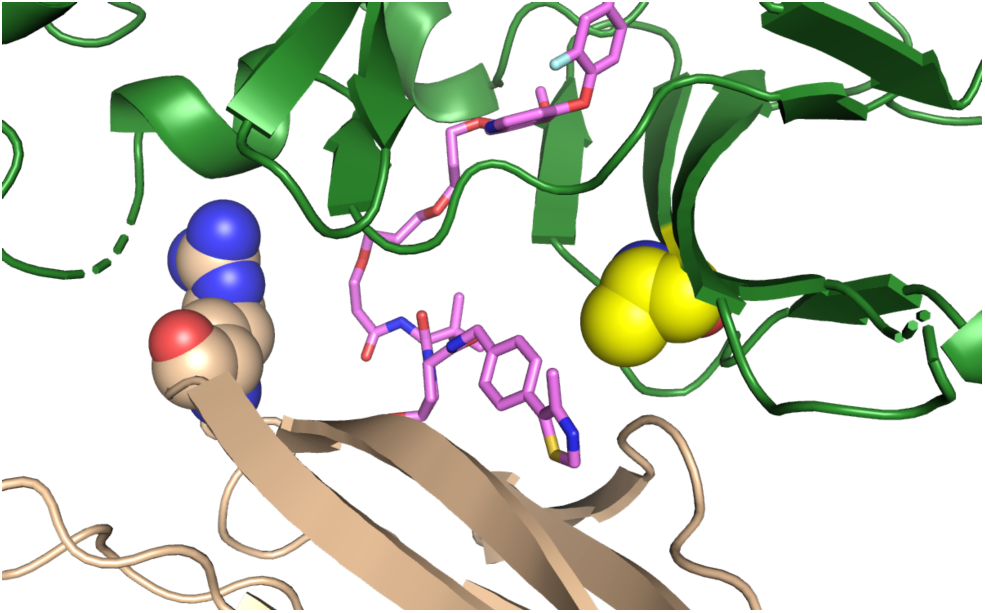
Model of the p38α/PROTAC1/VHL ternary complex. The lowest-energy model from the ensemble of the p38α/PROTAC1/VHL ternary complexes is shown. p38α is shown in *green*, PROTAC1 is *pink*, and VHL is *wheat*. Ala40 on p38α is shown in *yellow spheres*; mutation of this position to lysine was previously shown to disrupt this ternary complex. In contrast to a previously proposed model, our lowest-energy model positions Ala40 on opposite side of the interface as VHL’s Arg69 (*spheres*).

In summary, then, we find that although the warhead engages the structurally-conserved ATP-binding site in the same manner, and the kinase fold is conserved, our models of the ternary complex are notably different from one another. Rather, the protein-protein interactions in the ternary complex arise from the less-conserved regions on the kinase surface, which differ even among evolutionarily-related kinases such as p38α and p38δ. This in turn leads to PROTAC selectivity that can differ from that of the warhead in surprising and unexpected ways. This emergent selectivity will naturally depend as well on the E3 ligase that is to be recruited, as different E3 ligases will inevitably engage these non-biological binding partners in different ways.

## Discussion

To enable the results presented in this study, we first develop and describe a computational approach for evaluating the geometric and energetic “fit” of a given linker with respect to a pre-built ensemble of docked binding modes for the target protein and the E3 ligase. This approach allows us to rapidly filter the ensemble for candidate binding modes that are compatible with the constraints of the linker, and to build models of the target protein / PROTAC / E3 ligase ternary complex. The computational tools as well as the models of the ternary complexes are both disseminated as part of this study, in order to facilitate adoption of these methods and further analyses of the results presented here (see *Methods*).

Given our strategy for building these models of the ternary complex, a natural means for evaluating the linker’s “fit” was to calculate the fraction of the original docked ensemble compatible with the linker (which we dub *FFC*). As a result, our method does not produce a single model of the ternary complex, but rather a set of favorable models that are all consistent with the linker and which all include favorable interactions. Initially, our adoption of this strategy was in response to previous suggestions that crystal structure of PROTAC ternary complexes could be very susceptible to artifacts from crystal packing [52]. Very quickly, however, we recognized that our modeling yielded many models with different binding modes yet comparable energetics, suggesting that adoption of multiple binding modes may reflect a common underlying physical reality for PROTAC ternary complexes. Accordingly, conclusions drawn from a single static model (such a crystal structure) may be misleading; we therefore eagerly await insights from cryo-electron microscopy, which is exceptionally well-positioned to yield definitive answers regarding the range of motion in PROTAC ternary complexes.

At the outset, we envisioned that FFC would serve as a good measure of ternary complex formation from a biochemical standpoint: in other words, it might well recapitulate the degree to which the purified components might associate *in vitro*. Because cellular degradation is more commonly probed, however, it was more straightforward to draw these data from the literature; we were therefore delighted and surprised to discover how well the results from our computational pipeline agreed with this ultimately more-relevant endpoint. Importantly, our computational results do not direct incorporate any knowledge of a PROTAC’s cell penetrance, efflux, kinetics of ternary complex formation, ubiquitination efficiency once the ternary complex is formed, or rate of proteasomal degradation of the ubiquitinated target protein. This implies that – at least in the cases we explore here – a PROTAC’s activity is determined primarily by its ability to induce ternary complex formation and not these other factors. While systematic studies of these additional factors are only now beginning to be carried out [58], results from another recent report also align with this perspective. Cell permeability was measured for chloroalkane-tagged components of a JQ1-based degrader, and was found to be extremely low for the complete PROTAC: below even the concentration needed for effective degradation of BRD4, implying that this PROTAC must rely heavily on turnover to achieve depletion of its target protein [64]. This suggests that, for this series at least, cell permeability may not be the primary limitation of PROTAC activity.

A standing unanswered question with respect to PROTAC activity has been the role of cooperativity in ternary complex formation: in different systems, effective degraders have been identified that proved to be cooperative [35,50,51], non-cooperative [30], or even anti-cooperative [30,52]. Positive cooperativity implies that the target protein’s binding to the PROTAC is more favorable when the E3 ligase is present: this can occur if the ternary complex includes favorable interactions between the two proteins. By starting with an ensemble of docked models for the protein components, our computational strategy implicitly neglects binding modes in which the protein-protein interaction does not contribute favorably to the ternary complex. Instead, our approach focuses on binding modes in which the two proteins already have some intrinsic affinity for one another, and are therefore likely to exhibit (positive) cooperativity. Conversely, binding modes in which the arrangement of the protein components are unfavorable – leading to anti-cooperativity – may not be present in our docked ensemble, and thus could be missed in our subsequent analysis.

In an anti-cooperative scenario, addition of the PROTAC will still recruit the E3 ligase to the target protein: anti-cooperativity simply means that a higher concentration of PROTAC will be needed, relative to a non-cooperative or cooperative PROTAC with the same warhead and the E3-recruiting moiety. In a sense, anti-cooperative PROTACs degrade their targets *despite* their unfavorable interactions in the ternary complex. While these can certainly have cellular activity, optimizing a PROTAC linker for maximal ternary complex formation can be conceptually equated with maximizing cooperativity in the ternary complex. Thus, while our computational method may underestimate cellular activity of anti-cooperative PROTACs, we anticipate that even in these cases this approach can be used to facilitate optimization of activity through design of linkers that promote cooperativity.

As highlighted through the systematic comparisons to published reports that we present here, we envision that this computational strategy will prove valuable in at least three different contexts with respect to PROTAC design. First, for a given choice of warhead and E3 ligase, a single docked ensemble can be generated and numerous candidate linkers evaluated: as demonstrated (**Table 1**), this strategy can help select suitable linkers and provide a path for optimization that reduces the current requirement for synthesis and characterization of numerous PROTAC variants. Capitalizing on the suggestion that moving towards more rigid linkers can lead to improved potency and selectivity (as exemplified by a recent example using a spirocyclic pyrimidine [65]), this structure-based method is also well-positioned to suggest inflexible replacements for the typical PEG and alkyl linkers used in early-stage studies. Beyond simply varying the length or composition of the linker, the same strategy can also be used to explore variations in how the linker is attached to the warhead (**Table S1**). Second, subtle differences in the underlying complementarity between protein surfaces can lead to certain E3 ligases being better suited for degradation of a particular target protein (**Table 2**): our method is well-positioned to capture these differences, and thus guide the choice of an E3 ligase likely to yield effective degraders of the target protein. Third, a PROTAC’s preference across a protein family can also be explored, as a means to understanding how selectivity with respect to target degradation may differ from that of the warhead on which the PROTAC is built (**Table 2**).

By exploring ternary complexes of PROTACs built upon the same kinase-binding warhead, already we find that surface differences on the conserved kinase fold can change the preferred binding modes for a given PROTAC. While these differences are observed in the context of two PROTACs’ surprising degradation selectivity, already the underlying differences in the kinase/E3 binding preferences are presumably captured in the initial docked ensembles, even before the PROTAC linker has been specified. Moving forward, we anticipate that these preferences in the docked ensembled can be captured prior to building any models of ternary complexes, to evaluate the potential for selectivity irrespective of a particular linker. Support for this concept draws from recently-reported PROTACs built on the dual CDK6/CDK4 inhibitor palbociclib [66]. The dual selectivity of palbociclib is unsurprising, given that there is complete sequence conservation between these two kinases at all amino acid positions that contact this ATP-competitive inhibitor. Whereas multiple studies have described numerous CRBN-recruiting dual-selective degraders and CDK6-selective degraders [67-70], none report CDK4-selective degraders; the only reported CDK4-selective degraders are built on a slightly different warhead, ribociclib [71]. Based on the insights from the present study, we therefore speculate that CRBN can engage palbociclib-bound CDK6 in at least two binding modes: one of these can also be used to bind CDK4, and the other cannot. We envision that *a priori* knowledge of these available binding modes will enable rational design of selective degraders from the outset, by identifying PROTAC linkers that use binding modes unique to the protein target of interest.

Finally, looking to the future, we also expect that providing structural insights into the ensemble of ternary complexes used by a given PROTAC may yield additional insights beyond those demonstrated in the present study, such as anticipating means by which resistance to PROTACs may emerge [72].

## Conclusions

We report here the development of a new computational pipeline for structure-based modeling of PROTAC-mediated ternary complexes. By analysis of the resulting collections of models, we demonstrate that it is possible to retrospectively explain relationships between linker length/composition and cellular activity. Further, these models can also rationalize the surprising observation that the target-selectivity of a given PROTAC is not simply transferred from the target-binding warhead: rather, interactions with the E3 ligase in the ternary complex can dramatically shift the selectivity of the PROTAC. Moving forward, we expect that this approach can certainly be used to facilitate design of new PROTACs: by computationally screening libraries of candidate linkers, we envision prioritizing a very small number of compounds for synthesis and characterization. Given the current landscape in this field, which relies on synthesis of extensive collections of PROTAC variants in order to identify those yielding some amount of target degradation, we are confident that this new approach will facilitate design of PROTACs addressing many diverse targets.

## Methods

### Availability of computational tools and ternary complex models

Docking and minimization steps were carried out using the Rosetta software suite; Rosetta is freely available for academic use (www.rosettacommons.org). Conformer generation was carried out using OMEGA version 2.5.1.4.

Computational tools for building PROTAC ternary models are implemented in python. These tools are publicly available (along with complete documentation and examples of their usage) on Github (https://github.com/karanicolaslab/PROTAC_ternary).

To encourage further examination and additional analyses, the collection of ternary complex models for each portion of these studies are available on Mendeley: the ZXH set of Brd4BD1 degraders (**Table 1**) (http://dx.doi.org/10.17632/54hp53tfmt.1), the 28-31 set of Brd4BD1 degraders (**Table S1**) (http://dx.doi.org/10.17632/62hf4cvb7n.1), and the foretinib-based kinase degraders (**Table 2**) (http://dx.doi.org/10.17632/gdwxf3fmvr.1).

### Computational approach

To model a given system, the two relevant protein/ligand complex structures were downloaded from the PDB and first minimized using the Rosetta software suite. The two minimized complex structures were then manually combined together using the PyMol software, such that the two bound ligands were oriented towards one another. This starting structure was then prepared for docking using Rosetta’s pre-pack command:

~~~
docking_prepack_protocol.linuxgccrelease -s POI_ligand1_E3ligase_ligand2.pdb
      -use_input_sc -extra_res_fa ligand1.params ligand2.params
~~~

Protein-protein docking was then carried out as follows:

~~~
docking_protocol.linuxgccelease -s POI_ligand1_E3ligase_ligand2_prepacked.pdb
       -nstruct 50000 -use_input_sc -spin -dock_pert 5 20 -partners XY_MN
       -ex1 -ex2aro -extra_res_fa ligand1.params ligand2_params
       -out:file:scorefile score.sc -score:docking_interface_score 1
~~~

Here, the flag “-partners XY_MN” flag specifies that the small-molecule ligands must move together with their paired proteins. The “-dock_pert” flag specifies the extent of initial perturbations to be used in docking, and thus determines the search space. Only top 10% of the docking decoys were advanced to the next steps, as determined by their interaction energies (I_sc in the Rosetta output).

For each linker, the desired construct (i.e., the linker capped on either end with “stubs” from the protein-recruiting moieties) was built as a SMILES string then used as input for conformer generation with OMEGA as follows:

~~~
oeomega classic -in linker_name.smi -out linker_name.oeb.gz -maxconfs 1000
~~~

We used a custom python script to evaluate the RMSD of the stubs in a given linker conformer relative to their location in a docked model. If the RMSD of the stubs passes a specified cutoff value, the script then builds and outputs the complete model of the ternary complex. This script was used as follows:

~~~
ternary_model_prediction.py -d decoy.pdb -l linker_conformer.pdb
       -da decoy_atoms_list.txt -la linker_atoms_list.txt -c 0.4
       -wd decoy_atom_delete.txt -ld linker_atom_delete.txt
       -t default -r rmsd.txt
~~~

Instructions for the usage of this script and a complete explanation for each flag is distributed with the script itself (on Github).

### Statistical analysis

To evaluate statistical significance of the observed results, the original docked ensemble (5000 models) was split into five groups that each comprised 1000 docked models. Geometrical and energetic clustering was separately carried out for each subset, and FFC was independently calculated for each subset. Results are reported as the mean of each group, along with standard error of the mean. Statistical significance of differences in FFC amongst individual pairs of PROTACs were computed using the (unpaired, two-tailed) Student’s t-test.

### PDB structures used in calculations

Studies of the “ZXH” series and the “28-31” series were initiated from the JQ1-bound crystal structure of Brd4^BD1^ (PDB 3mxf) and the lenalidomide-bound crystal structure of CRBN (PDB 4tz4).

Studies of the foretinib-based PROTACs were initiated from the foretinib-bound crystal structures of c-Met (PDB 3lq8), EphA2 (PDB 5ia4), and STK10 (PDB 6i2y). These were each paired with crystal structures of VHL (PDB 4w9h) and CRBN (PDB 4tz4) bound to their respective ligands. Because crystal structures were not available for foretinib bound to p38α or p38δ, these were generated via comparative modeling. The complete protocol for building these models is described in the context of a separate study [73].

## Acknowledgements

We thank Sven Miller for valuable feedback on the documentation for this code in preparation for its dissemination. This work used the Extreme Science and Engineering Discovery Environment (XSEDE) allocation MCB130049, which is supported by National Science Foundation grant number ACI-1548562. This work was supported by grants from the National Institute of General Medical Sciences (R01GM123336) and from the National Science Foundation (CHE-1836950). This research was funded in part through the NIH/NCI Cancer Center Support Grant P30 CA006927.

## Notes

### Competing Interest Statement

The authors have declared no competing interest.

## References

1. Sakamoto KM, Kim KB, Kumagai A, Mercurio F, Crews CM, Deshaies RJ. Protacs: chimeric molecules that target proteins to the Skp1-Cullin-F box complex for ubiquitination and degradation. Proc Natl Acad Sci U S A. 2001; 98:8554–9.

2. Metzger MB, Pruneda JN, Klevit RE, Weissman AM. RING-type E3 ligases: master manipulators of E2 ubiquitin-conjugating enzymes and ubiquitination. Biochim Biophys Acta. 2014; 1843:47–60.

3. Barry M, Fruh K. Viral modulators of cullin RING ubiquitin ligases: culling the host defense. Sci STKE. 2006; 2006:pe21.

4. Ito T, Ando H, Suzuki T, Ogura T, Hotta K, Imamura Y, Yamaguchi Y, Handa H. Identification of a primary target of thalidomide teratogenicity. Science. 2010; 327:1345–50.

5. Donovan KA, An J, Nowak RP, Yuan JC, Fink EC, Berry BC, Ebert BL, Fischer ES. Thalidomide promotes degradation of SALL4, a transcription factor implicated in Duane Radial Ray syndrome. Elife. 2018; 7.

6. Asatsuma-Okumura T, Ando H, De Simone M, Yamamoto J, Sato T, Shimizu N, Asakawa K, Yamaguchi Y, Ito T, Guerrini L, Handa H. p63 is a cereblon substrate involved in thalidomide teratogenicity. Nat Chem Biol. 2019; 15:1077–84.

7. Fischer ES, Bohm K, Lydeard JR, Yang H, Stadler MB, Cavadini S, Nagel J, Serluca F, Acker V, Lingaraju GM, Tichkule RB, Schebesta M, Forrester WC, Schirle M, Hassiepen U, Ottl J, Hild M, Beckwith RE, Harper JW, Jenkins JL, Thoma NH. Structure of the DDB1-CRBN E3 ubiquitin ligase in complex with thalidomide. Nature. 2014; 512:49–53.

8. Neklesa TK, Winkler JD, Crews CM. Targeted protein degradation by PROTACs. Pharmacol Ther. 2017; 174:138–44.

9. Bondeson DP, Mares A, Smith IE, Ko E, Campos S, Miah AH, Mulholland KE, Routly N, Buckley DL, Gustafson JL, Zinn N, Grandi P, Shimamura S, Bergamini G, Faelth-Savitski M, Bantscheff M, Cox C, Gordon DA, Willard RR, Flanagan JJ, Casillas LN, Votta BJ, den Besten W, Famm K, Kruidenier L, Carter PS, Harling JD, Churcher I, Crews CM. Catalytic in vivo protein knockdown by small-molecule PROTACs. Nat Chem Biol. 2015; 11:611–7.

10. Cyrus K, Wehenkel M, Choi EY, Lee H, Swanson H, Kim KB. Jostling for position: optimizing linker location in the design of estrogen receptor-targeting PROTACs. ChemMedChem. 2010; 5:979–85.

11. Wang L, Guillen VS, Sharma N, Flessa K, Min J, Carlson KE, Toy W, Braqi S, Katzenellenbogen BS, Katzenellenbogen JA, Chandarlapaty S, Sharma A. New Class of Selective Estrogen Receptor Degraders (SERDs): Expanding the Toolbox of PROTAC Degrons. ACS Med Chem Lett. 2018; 9:803–8.

12. Sakamoto KM, Kim KB, Verma R, Ransick A, Stein B, Crews CM, Deshaies RJ. Development of Protacs to target cancer-promoting proteins for ubiquitination and degradation. Mol Cell Proteomics. 2003; 2:1350–8.

13. Rodriguez-Gonzalez A, Cyrus K, Salcius M, Kim K, Crews CM, Deshaies RJ, Sakamoto KM. Targeting steroid hormone receptors for ubiquitination and degradation in breast and prostate cancer. Oncogene. 2008; 27:7201–11.

14. Winter GE, Buckley DL, Paulk J, Roberts JM, Souza A, Dhe-Paganon S, Bradner JE. Phthalimide conjugation as a strategy for in vivo target protein degradation. Science. 2015; 348:1376–81.

15. Lu J, Qian Y, Altieri M, Dong H, Wang J, Raina K, Hines J, Winkler JD, Crew AP, Coleman K, Crews CM. Hijacking the E3 Ubiquitin Ligase Cereblon to Efficiently Target BRD4. Chem Biol. 2015; 22:755–63.

16. Zengerle M, Chan K-H, Ciulli A. Selective small molecule induced degradation of the BET bromodomain protein BRD4. ACS chemical biology. 2015; 10:1770–7.

17. Lai AC, Toure M, Hellerschmied D, Salami J, Jaime-Figueroa S, Ko E, Hines J, Crews CM. Modular PROTAC Design for the Degradation of Oncogenic BCR-ABL. Angew Chem Int Ed Engl. 2016; 55:807–10.

18. Robb CM, Contreras JI, Kour S, Taylor MA, Abid M, Sonawane YA, Zahid M, Murry DJ, Natarajan A, Rana S. Chemically induced degradation of CDK9 by a proteolysis targeting chimera (PROTAC). Chem Commun (Camb). 2017; 53:7577–80.

19. Burslem GM, Smith BE, Lai AC, Jaime-Figueroa S, McQuaid DC, Bondeson DP, Toure M, Dong H, Qian Y, Wang J, Crew AP, Hines J, Crews CM. The Advantages of Targeted Protein Degradation Over Inhibition: An RTK Case Study. Cell Chem Biol. 2018; 25:67–77 e3.

20. Bondeson DP, Smith BE, Burslem GM, Buhimschi AD, Hines J, Jaime-Figueroa S, Wang J, Hamman BD, Ishchenko A, Crews CM. Lessons in PROTAC Design from Selective Degradation with a Promiscuous Warhead. Cell Chem Biol. 2018; 25:78–87 e5.

21. Zhang C, Han XR, Yang X, Jiang B, Liu J, Xiong Y, Jin J. Proteolysis Targeting Chimeras (PROTACs) of Anaplastic Lymphoma Kinase (ALK). Eur J Med Chem. 2018; 151:304–14.

22. Crew AP, Raina K, Dong H, Qian Y, Wang J, Vigil D, Serebrenik YV, Hamman BD, Morgan A, Ferraro C, Siu K, Neklesa TK, Winkler JD, Coleman KG, Crews CM. Identification and Characterization of Von Hippel-Lindau-Recruiting Proteolysis Targeting Chimeras (PROTACs) of TANK-Binding Kinase 1. J Med Chem. 2018; 61:583–98.

23. Huang HT, Dobrovolsky D, Paulk J, Yang G, Weisberg EL, Doctor ZM, Buckley DL, Cho JH, Ko E, Jang J, Shi K, Choi HG, Griffin JD, Li Y, Treon SP, Fischer ES, Bradner JE, Tan L, Gray NS. A Chemoproteomic Approach to Query the Degradable Kinome Using a Multi-kinase Degrader. Cell Chem Biol. 2018; 25:88–99 e6.

24. Olson CM, Jiang B, Erb MA, Liang Y, Doctor ZM, Zhang Z, Zhang T, Kwiatkowski N, Boukhali M, Green JL, Haas W, Nomanbhoy T, Fischer ES, Young RA, Bradner JE, Winter GE, Gray NS. Pharmacological perturbation of CDK9 using selective CDK9 inhibition or degradation. Nat Chem Biol. 2018; 14:163–70.

25. Powell CE, Gao Y, Tan L, Donovan KA, Nowak RP, Loehr A, Bahcall M, Fischer ES, Janne PA, George RE, Gray NS. Chemically Induced Degradation of Anaplastic Lymphoma Kinase (ALK). J Med Chem. 2018; 61:4249–55.

26. Cromm PM, Samarasinghe KTG, Hines J, Crews CM. Addressing Kinase-Independent Functions of Fak via PROTAC-Mediated Degradation. J Am Chem Soc. 2018; 140:17019–26.

27. Chen H, Chen F, Liu N, Wang X, Gou S. Chemically induced degradation of CK2 by proteolysis targeting chimeras based on a ubiquitin-proteasome pathway. Bioorg Chem. 2018; 81:536–44.

28. Buhimschi AD, Armstrong HA, Toure M, Jaime-Figueroa S, Chen TL, Lehman AM, Woyach JA, Johnson AJ, Byrd JC, Crews CM. Targeting the C481S Ibrutinib-Resistance Mutation in Bruton’s Tyrosine Kinase Using PROTAC-Mediated Degradation. Biochemistry. 2018; 57:3564–75.

29. Sun Y, Zhao X, Ding N, Gao H, Wu Y, Yang Y, Zhao M, Hwang J, Song Y, Liu W, Rao Y. PROTAC-induced BTK degradation as a novel therapy for mutated BTK C481S induced ibrutinib-resistant B-cell malignancies. Cell Res. 2018; 28:779–81.

30. Zorba A, Nguyen C, Xu Y, Starr J, Borzilleri K, Smith J, Zhu H, Farley KA, Ding W, Schiemer J, Feng X, Chang JS, Uccello DP, Young JA, Garcia-Irrizary CN, Czabaniuk L, Schuff B, Oliver R, Montgomery J, Hayward MM, Coe J, Chen J, Niosi M, Luthra S, Shah JC, El-Kattan A, Qiu X, West GM, Noe MC, Shanmugasundaram V, Gilbert AM, Brown MF, Calabrese MF. Delineating the role of cooperativity in the design of potent PROTACs for BTK. Proc Natl Acad Sci U S A. 2018; 115:E7285–E92.

31. Bian J, Ren J, Li Y, Wang J, Xu X, Feng Y, Tang H, Wang Y, Li Z. Discovery of Wogonin-based PROTACs against CDK9 and capable of achieving antitumor activity. Bioorg Chem. 2018; 81:373–81.

32. Kang CH, Lee DH, Lee CO, Du Ha J, Park CH, Hwang JY. Induced protein degradation of anaplastic lymphoma kinase (ALK) by proteolysis targeting chimera (PROTAC). Biochem Biophys Res Commun. 2018; 505:542–7.

33. Burslem GM, Song J, Chen X, Hines J, Crews CM. Enhancing Antiproliferative Activity and Selectivity of a FLT-3 Inhibitor by Proteolysis Targeting Chimera Conversion. J Am Chem Soc. 2018; 140:16428–32.

34. Gechijian LN, Buckley DL, Lawlor MA, Reyes JM, Paulk J, Ott CJ, Winter GE, Erb MA, Scott TG, Xu M, Seo HS, Dhe-Paganon S, Kwiatkowski NP, Perry JA, Qi J, Gray NS, Bradner JE. Functional TRIM24 degrader via conjugation of ineffectual bromodomain and VHL ligands. Nat Chem Biol. 2018; 14:405–12.

35. Farnaby W, Koegl M, Roy MJ, Whitworth C, Diers E, Trainor N, Zollman D, Steurer S, Karolyi-Oezguer J, Riedmueller C, Gmaschitz T, Wachter J, Dank C, Galant M, Sharps B, Rumpel K, Traxler E, Gerstberger T, Schnitzer R, Petermann O, Greb P, Weinstabl H, Bader G, Zoephel A, Weiss-Puxbaum A, Ehrenhofer-Wolfer K, Wohrle S, Boehmelt G, Rinnenthal J, Arnhof H, Wiechens N, Wu MY, Owen-Hughes T, Ettmayer P, Pearson M, McConnell DB, Ciulli A. BAF complex vulnerabilities in cancer demonstrated via structure-based PROTAC design. Nat Chem Biol. 2019; 15:672–80.

36. Chu TT, Gao N, Li QQ, Chen PG, Yang XF, Chen YX, Zhao YF, Li YM. Specific Knockdown of Endogenous Tau Protein by Peptide-Directed Ubiquitin-Proteasome Degradation. Cell Chem Biol. 2016; 23:453–61.

37. Lu M, Liu T, Jiao Q, Ji J, Tao M, Liu Y, You Q, Jiang Z. Discovery of a Keap1-dependent peptide PROTAC to knockdown Tau by ubiquitination-proteasome degradation pathway. Eur J Med Chem. 2018; 146:251–9.

38. Buckley DL, Raina K, Darricarrere N, Hines J, Gustafson JL, Smith IE, Miah AH, Harling JD, Crews CM. HaloPROTACS: Use of Small Molecule PROTACs to Induce Degradation of HaloTag Fusion Proteins. ACS Chem Biol. 2015; 10:1831–7.

39. Zhao Q, Lan T, Su S, Rao Y. Induction of apoptosis in MDA-MB-231 breast cancer cells by a PARP1-targeting PROTAC small molecule. Chem Commun (Camb). 2019; 55:369–72.

40. Tinworth CP, Lithgow H, Dittus L, Bassi ZI, Hughes SE, Muelbaier M, Dai H, Smith IED, Kerr WJ, Burley GA, Bantscheff M, Harling JD. PROTAC-Mediated Degradation of Bruton’s Tyrosine Kinase Is Inhibited by Covalent Binding. ACS Chem Biol. 2019; 14:342–7.

41. Ward CC, Kleinman JI, Brittain SM, Lee PS, Chung CYS, Kim K, Petri Y, Thomas JR, Tallarico JA, McKenna JM, Schirle M, Nomura DK. Covalent Ligand Screening Uncovers a RNF4 E3 Ligase Recruiter for Targeted Protein Degradation Applications. ACS Chem Biol. 2019; 14:2430–40.

42. Ottis P, Toure M, Cromm PM, Ko E, Gustafson JL, Crews CM. Assessing Different E3 Ligases for Small Molecule Induced Protein Ubiquitination and Degradation. ACS Chem Biol. 2017; 12:2570–8.

43. Zhang X, Crowley VM, Wucherpfennig TG, Dix MM, Cravatt BF. Electrophilic PROTACs that degrade nuclear proteins by engaging DCAF16. Nat Chem Biol. 2019; 15:737–46.

44. Cyrus K, Wehenkel M, Choi EY, Han HJ, Lee H, Swanson H, Kim KB. Impact of linker length on the activity of PROTACs. Mol Biosyst. 2011; 7:359–64.

45. Steinebach C, Sosic I, Lindner S, Bricelj A, Kohl F, Ng YLD, Monschke M, Wagner KG, Kronke J, Gutschow M. A MedChem toolbox for cereblon-directed PROTACs. Medchemcomm. 2019; 10:1037–41.

46. Zoppi V, Hughes SJ, Maniaci C, Testa A, Gmaschitz T, Wieshofer C, Koegl M, Riching KM, Daniels DL, Spallarossa A, Ciulli A. Iterative Design and Optimization of Initially Inactive Proteolysis Targeting Chimeras (PROTACs) Identify VZ185 as a Potent, Fast, and Selective von Hippel-Lindau (VHL) Based Dual Degrader Probe of BRD9 and BRD7. J Med Chem. 2019; 62:699–726.

47. Chamberlain PP, Lopez-Girona A, Miller K, Carmel G, Pagarigan B, Chie-Leon B, Rychak E, Corral LG, Ren YJ, Wang M, Riley M, Delker SL, Ito T, Ando H, Mori T, Hirano Y, Handa H, Hakoshima T, Daniel TO, Cathers BE. Structure of the human Cereblon-DDB1-lenalidomide complex reveals basis for responsiveness to thalidomide analogs. Nat Struct Mol Biol. 2014; 21:803–9.

48. Galdeano C, Gadd MS, Soares P, Scaffidi S, Van Molle I, Birced I, Hewitt S, Dias DM, Ciulli A. Structure-guided design and optimization of small molecules targeting the protein-protein interaction between the von Hippel-Lindau (VHL) E3 ubiquitin ligase and the hypoxia inducible factor (HIF) alpha subunit with in vitro nanomolar affinities. J Med Chem. 2014; 57:8657–63.

49. Drummond ML, Williams CI. In Silico Modeling of PROTAC-Mediated Ternary Complexes: Validation and Application. J Chem Inf Model. 2019; 59:1634–44.

50. Gadd MS, Testa A, Lucas X, Chan KH, Chen W, Lamont DJ, Zengerle M, Ciulli A. Structural basis of PROTAC cooperative recognition for selective protein degradation. Nat Chem Biol. 2017; 13:514–21.

51. Roy MJ, Winkler S, Hughes SJ, Whitworth C, Galant M, Farnaby W, Rumpel K, Ciulli A. SPR-Measured Dissociation Kinetics of PROTAC Ternary Complexes Influence Target Degradation Rate. ACS Chem Biol. 2019; 14:361–8.

52. Nowak RP, DeAngelo SL, Buckley D, He Z, Donovan KA, An J, Safaee N, Jedrychowski MP, Ponthier CM, Ishoey M, Zhang T, Mancias JD, Gray NS, Bradner JE, Fischer ES. Plasticity in binding confers selectivity in ligand-induced protein degradation. Nat Chem Biol. 2018; 14:706–14.

53. Leaver-Fay A, Tyka M, Lewis SM, Lange OF, Thompson J, Jacak R, Kaufman K, Renfrew PD, Smith CA, Sheffler W, Davis IW, Cooper S, Treuille A, Mandell DJ, Richter F, Ban YE, Fleishman SJ, Corn JE, Kim DE, Lyskov S, Berrondo M, Mentzer S, Popovic Z, Havranek JJ, Karanicolas J, Das R, Meiler J, Kortemme T, Gray JJ, Kuhlman B, Baker D, Bradley P. ROSETTA3: an object-oriented software suite for the simulation and design of macromolecules. Methods Enzymol. 2011; 487:545–74.

54. Weitzner BD, Jeliazkov JR, Lyskov S, Marze N, Kuroda D, Frick R, Adolf-Bryfogle J, Biswas N, Dunbrack RL, Jr., Gray JJ. Modeling and docking of antibody structures with Rosetta. Nat Protoc. 2017; 12:401–16.

55. Hawkins PC, Skillman AG, Nicholls A. Comparison of shape-matching and docking as virtual screening tools. J Med Chem. 2007; 50:74–82.

56. Hawkins PC, Nicholls A. Conformer generation with OMEGA: learning from the data set and the analysis of failures. J Chem Inf Model. 2012; 52:2919–36.

57. Alford RF, Leaver-Fay A, Jeliazkov JR, O’Meara MJ, DiMaio FP, Park H, Shapovalov MV, Renfrew PD, Mulligan VK, Kappel K, Labonte JW, Pacella MS, Bonneau R, Bradley P, Dunbrack RL, Jr., Das R, Baker D, Kuhlman B, Kortemme T, Gray JJ. The Rosetta All-Atom Energy Function for Macromolecular Modeling and Design. J Chem Theory Comput. 2017; 13:3031–48.

58. Hughes SJ, Ciulli A. Molecular recognition of ternary complexes: a new dimension in the structure-guided design of chemical degraders. Essays Biochem. 2017; 61:505–16.

59. Riching KM, Mahan S, Corona CR, McDougall M, Vasta JD, Robers MB, Urh M, Daniels DL. Quantitative Live-Cell Kinetic Degradation and Mechanistic Profiling of PROTAC Mode of Action. ACS Chem Biol. 2018; 13:2758–70.

60. Qin C, Hu Y, Zhou B, Fernandez-Salas E, Yang CY, Liu L, McEachern D, Przybranowski S, Wang M, Stuckey J, Meagher J, Bai L, Chen Z, Lin M, Yang J, Ziazadeh DN, Xu F, Hu J, Xiang W, Huang L, Li S, Wen B, Sun D, Wang S. Discovery of QCA570 as an Exceptionally Potent and Efficacious Proteolysis Targeting Chimera (PROTAC) Degrader of the Bromodomain and Extra-Terminal (BET) Proteins Capable of Inducing Complete and Durable Tumor Regression. J Med Chem. 2018; 61:6685–704.

61. Filippakopoulos P, Qi J, Picaud S, Shen Y, Smith WB, Fedorov O, Morse EM, Keates T, Hickman TT, Felletar I, Philpott M, Munro S, McKeown MR, Wang Y, Christie AL, West N, Cameron MJ, Schwartz B, Heightman TD, La Thangue N, French CA, Wiest O, Kung AL, Knapp S, Bradner JE. Selective inhibition of BET bromodomains. Nature. 2010; 468:1067–73.

62. Qian F, Engst S, Yamaguchi K, Yu P, Won KA, Mock L, Lou T, Tan J, Li C, Tam D, Lougheed J, Yakes FM, Bentzien F, Xu W, Zaks T, Wooster R, Greshock J, Joly AH. Inhibition of tumor cell growth, invasion, and metastasis by EXEL-2880 (XL880, GSK1363089), a novel inhibitor of HGF and VEGF receptor tyrosine kinases. Cancer Res. 2009; 69:8009–16.

63. Heinzlmeir S, Kudlinzki D, Sreeramulu S, Klaeger S, Gande SL, Linhard V, Wilhelm M, Qiao H, Helm D, Ruprecht B, Saxena K, Medard G, Schwalbe H, Kuster B. Chemical Proteomics and Structural Biology Define EPHA2 Inhibition by Clinical Kinase Drugs. ACS Chem Biol. 2016; 11:3400–11.

64. Foley CA, Potjewyd F, Lamb KN, James LI, Frye SV. Assessing the Cell Permeability of Bivalent Chemical Degraders Using the Chloroalkane Penetration Assay. ACS Chem Biol. 2020; 15:290–5.

65. Nunes J, McGonagle GA, Eden J, Kiritharan G, Touzet M, Lewell X, Emery J, Eidam H, Harling JD, Anderson NA. Targeting IRAK4 for Degradation with PROTACs. ACS Med Chem Lett. 2019; 10:1081–5.

66. Fry DW, Harvey PJ, Keller PR, Elliott WL, Meade M, Trachet E, Albassam M, Zheng X, Leopold WR, Pryer NK, Toogood PL. Specific inhibition of cyclin-dependent kinase 4/6 by PD 0332991 and associated antitumor activity in human tumor xenografts. Mol Cancer Ther. 2004; 3:1427–38.

67. Brand M, Jiang B, Bauer S, Donovan KA, Liang Y, Wang ES, Nowak RP, Yuan JC, Zhang T, Kwiatkowski N, Muller AC, Fischer ES, Gray NS, Winter GE. Homolog-Selective Degradation as a Strategy to Probe the Function of CDK6 in AML. Cell Chem Biol. 2019; 26:300–6 e9.

68. Anderson NA, Cryan J, Ahmed A, Dai H, McGonagle GA, Rozier C, Benowitz AB. Selective CDK6 degradation mediated by cereblon, VHL, and novel IAP-recruiting PROTACs. Bioorg Med Chem Lett. 2020; 30:127106.

69. Steinebach C, Ng YLD, Sosič I, Lee C-S, Chen S, Lindner S, Vu LP, Bricelj A, Haschemi R, Monschke M, Steinwarz E, Wagner KG, Bendas G, Luo J, Gütschow M, Krönke J. Systematic exploration of different E3 ubiquitin ligases: an approach towards potent and selective CDK6 degraders. Chemical Science. 2020; 11:3474–86.

70. Rana S, Bendjennat M, Kour S, King HM, Kizhake S, Zahid M, Natarajan A. Selective degradation of CDK6 by a palbociclib based PROTAC. Bioorg Med Chem Lett. 2019; 29:1375–9.

71. Jiang B, Wang ES, Donovan KA, Liang Y, Fischer ES, Zhang T, Gray NS. Development of Dual and Selective Degraders of Cyclin-Dependent Kinases 4 and 6. Angew Chem Int Ed Engl. 2019; 58:6321–6.

72. Zhang L, Riley-Gillis B, Vijay P, Shen Y. Acquired Resistance to BET-PROTACs (Proteolysis-Targeting Chimeras) Caused by Genomic Alterations in Core Components of E3 Ligase Complexes. Mol Cancer Ther. 2019; 18:1302–11.

73. Kirubakaran P, Morton G, Zhang P, Zhang H, Gordon J, Abou-Gharbia M, Issa J-PJ, Wu J, Childers W, Karanicolas J. Comparative Modeling of CDK9 Inhibitors to Explore Selectivity and Structure-Activity Relationships. submitted.

